# A High-throughput Screening Method to Identify Proteins Involved in Unfolded Protein Response Signaling in Plants

**DOI:** 10.1101/825190

**Authors:** André Alcântara, Denise Seitner, Fernando Navarrete, Armin Djamei

**Affiliations:** Gregor Mendel Institute of Molecular Plant Biology, Vienna, Austria; Leibniz-Institut für Pflanzengenetik und Kulturpflanzenforschung (IPK), Gatersleben, Germany

**Keywords:** Unfolded protein response (UPR), high-throughput, *Nicotiana benthamiana*, transient expression, *Ustilago maydis*

## Abstract

**Background:** The unfolded protein response (UPR) is a highly conserved process in eukaryotic organisms that plays a crucial role in adaptation and development. While the most ubiquitous components of this pathway have been characterized, current efforts are focused on identifying and characterizing other UPR factors that play a role in specific conditions, such as developmental changes, abiotic cues, and biotic interactions. Considering the central role of protein secretion in plant pathogen interactions, there has also been a recent focus on understanding how pathogens manipulate their host’s UPR to facilitate infection.

**Results:** We developed a high-throughput screening assay to identify proteins that interfere with UPR signaling *in planta*. A set of 35 genes from a library of secreted proteins from the maize pathogen *Ustilago maydis* were transiently co-expressed with a reporter construct that upregulates enhanced yellow fluorescent protein (eYFP) expression upon UPR stress in *Nicotiana benthamiana* plants. After UPR stress induction, leaf discs were placed in 96 well plates and eYFP expression was measured. This allowed us to identify a previously undescribed fungal protein that inhibits plant UPR signaling, which was then confirmed using the classical but more laborious qRT-PCR method.

**Conclusions:** We have established a rapid and reliable fluorescence-based method to identify heterologously expressed proteins involved in UPR stress in plants. This system can be used for initial screens with libraries of proteins and potentially other molecules to identify candidates for further validation and characterization.

## Background

The unfolded protein response (UPR) is a conserved mechanism across eukaryotic organisms for maintaining homeostasis in the endoplasmic reticulum (ER). Proteins from the secretory pathway are translated into the ER where they acquire their native folding and undergo posttranslational modifications. Then, the proteins are shuttled to other organelles, for further processing, or directly to their target compartment, to fulfil their functions. Due to their sessile nature, plants rely heavily on the secretory pathway to respond to changes in, and interact with, their environment. A change in environmental stimuli can lead to significant changes in a cell’s transcriptional programing, which in turn cause an overloading of the ER with newly synthesized proteins. These overwhelm the chaperones within it, leading to the accumulation of unfolded proteins, which causes ER stress (Chakraborty et al., 2016; Nawkar et al., 2018; Strasser, 2018). Examples of environmental factors that can lead to UPR include temperature changes, ionic and osmotic stresses, high light, heavy metal toxicity, and biotic interactions (Gao et al., 2008; Liu et al., 2007; Meng et al., 2017; Moreno et al., 2012; Nawkar et al., 2017; Valente et al., 2009; Zhang et al., 2019). Together with changes in developmental programming, these deviations from cellular homeostasis can lead to protein oxidation and/or defects in protein glycosylation that lead to their denaturation and accumulation in different organelles, including the ER, leading to stress.

In plants, there are at least two different mechanisms by which ER stress can be perceived and activate a signaling cascade that triggers UPR. In the Inositol-requiring enzyme 1 (IRE1) pathway, luminal binding proteins (BiPs) interact with the ER-membrane protein IRE1 in the ER lumen. When unfolded proteins accumulate, they are bound by BiPs, releasing IRE1 proteins that then form dimers which unconventionally splice basic leucin zipper (bZip) 60 mRNAs in the cytosol. The spliced mRNA translates into a functional transcription factor that shuttles to the nucleus and promotes the upregulation of genes that contain UPR responsive elements (UPREs) and ER stress elements (ERSEs) in their regulatory regions (Hayashi et al., 2013; Mori et al., 1996; Sun et al., 2013). The other UPR signaling pathway involves the ER-membrane bZips 17 and 28, which are also bound by BiPs. Upon their release, they are transported to the Golgi apparatus. There, two proteases cleave the full length protein – the site 1 protease (S1P) in the C-terminal region inside the Golgi and the site 2 protease (S2P) in the cytosol – releasing the transcription factor which then migrates to the nucleus and upregulates ER stress genes (Gao et al., 2008). Both signaling pathways ultimately lead to the upregulation of genes to either correctly fold or degrade misfolded proteins, and to regulate transcription and translation to restore ER homeostasis (Iwata et al., 2010a; Srivastava et al., 2018). Transient ER stress can be relieved by the UPR, while persistent ER stress may lead to programmed cell death (PCD; Moreno & Orellana, 2011).

Some of the downstream targets of UPR signaling include genes related to plant immunity. Biotic stresses cause dramatic changes in the host’s transcriptional programing that lead to UPR (Moreno et al., 2012; Xu et al., 2019). Depending on their lifestyle, plant pathogens evolved mechanisms to either promote PCD – in the case of necrotrophic organisms – or to inhibit it and other immune responses – in the case of biotrophs. It is therefore not surprising that plant UPR components were recently reported as targets of the molecules pathogens secrete to control their host (i.e. effectors). For instance, after determining that the *Phytophthora sojae* effector Avh262 was required for full virulence, Jing et al. (2016) transiently expressed it in *N. benthamiana* fused to a green fluorescent protein. Co-immunoprecipitation followed by mass spectrometry revealed that PsAvh262 binds to BiP proteins and further experiments showed that stabilization of this target dampens plant resistance. More recently, the *Phytophthora capsica* effector Avr3a12 was found to interact with FKBP15-2, a plant peptidyl-prolyl cis-trans isomerase which was found to be required for ER stress mediated immunity (Fan et al., 2018). However, the lack of a method for screening proteins that interfere with plant UPR has made it difficult to identify effectors in other pathogens that might play a role in this process.

Though the conserved pathways of UPR signaling in plants have been described, a number of factors involved in its regulation remain to be characterized. Due to its central role in various stress responses, methods for identifying UPR modulators in specific conditions are crucial to advance our understanding of this cellular mechanism. Chen & Brandizzi (2013) described different ways of inducing ER stress in *Arabidopsis thaliana* plants and measure their effects through quantitative polymerase chain reaction (qPCR) measurement of UPR target genes. Another method was described by McCormack et al. (2015) who developed a screening assay to test the sensitivity of *A. thaliana* seedlings to tunicamycin (Tm) – an N-glycosylation inhibitor that causes ER stress and triggers UPR – in response to different stimuli and/or with different genetic backgrounds. Additionally, other authors have adapted protocols to investigate the specific role of their proteins of interest in UPR (Hayashi et al., 2013; Liu & Howell, 2010; Meng et al., 2017; Nawkar et al., 2017) but a simple, reliable, high-throughput method to identify new proteins, and potentially other small molecules or environmental conditions, involved in this mechanism is yet to be reported.

Here we report a method for screening proteins, and potentially other molecules or conditions, that influence plant UPR. This method relies on fluorescence measurements of *Nicotiana benthamiana* leaf discs transiently expressing two genetic constructs. One of them expresses the protein of interest, while the second plasmid encodes an ER-stress responsive promoter controlling the expression of enhanced yellow fluorescent protein (eYFP). By using a subset of proteins from a library of secreted proteins (*i.e.* putative effectors) from the maize pathogen *Ustilago maydis*, we were able to identify one protein that inhibits UPR signaling in plants. After validation by more classical, laborious methods, this simple approach allows for the screening and identification of new players in plant UPR that may have a role in specific conditions.

## Results

### A fluorescence-based assay to measure UPR stress

We developed a method that measures relative UPR stress and signaling in plants (Fig. 1). By co-expressing a reporter construct and a protein of interest, interference in UPR signaling can be assessed and new players in this cellular mechanism can be identified. Additionally, the same reporter plasmid could be used to assess the influence of other molecules or environmental conditions on UPR signaling.

**Fig. 1.**
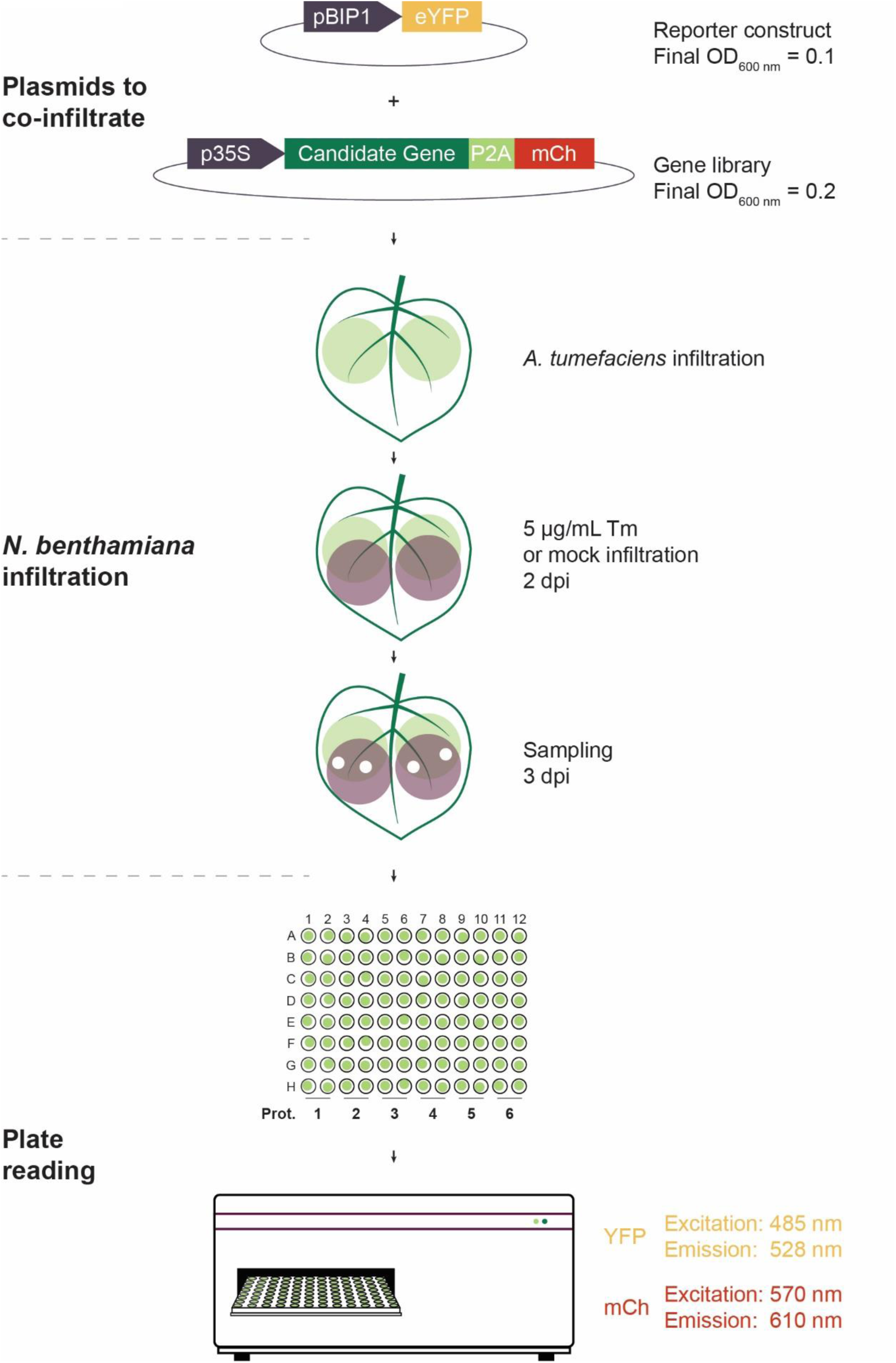
Graphical protocol to screen proteins for influence on unfolded protein response (UPR) signaling. Candidate genes are cloned into a plant expression vector and co-infiltrated with the reporter plasmid at different 600 nm optical densities (OD_600 nm_), into *N. benthamiana* leaves. Two days post-infiltration (dpi), the same leaves are infiltrated with either tunicamycin (Tm) or DMSO (mock) to assess inhibition or induction of UPR signaling, respectively. At 3 dpi, leaf discs are sampled and floated on water in 96 well plates. Fluorescence intensity is measured in a plate reader. pBIP1 – regulatory region of the BiP1 protein from *A. thaliana*; eYFP – enhanced Yellow Fluorescent Protein; mCh – mCherry; p35S – CaMV 35S promoter; P2A -porcine teschovirus-1 2A “self-cleaving” peptide

In brief, candidate genes are cloned in an expression vector under the control of the CaMV 35S promoter (p35S). An mCherry (mCh) fluorophore coding sequence is cloned in frame with the candidate gene but is separated by the porcine teschovirus-1 2A (P2A) self-cleaving peptide (Kim et al., 2011). This results in strong expression of the proteins of interest with a small C-terminal tag – which minimizes interference with the native folding and function – and the separate expression of a fluorophore in equimolar amounts. mCh fluorescence is then used as a proxy for transformation efficiency and relative protein expression levels. A library of constructs with proteins of interest can easily be generated to efficiently test for UPR interference. A construct with a second mCh coding sequence instead of the gene of interest is used as a reference (*i.e.* a construct that does not interfere with UPR signaling). Each construct is electroporated into *Agrobacterium tumefaciens* strains and co-infiltrated in *N. benthamiana* plants with a reporter construct expressing eYFP under the control of the ER stress inducible promoter pBIP1. Two days after infiltration, the same *N. benthamiana* leaves are infiltrated with either 0.5% DMSO, as a mock treatment, or 5 μg/mL of tunicamycin (Tm), to induce ER stress and UPR signaling. Approximately 24 hours after the second infiltration, leaf discs are sampled and floated on water in 96 well plates. eYFP and mCh fluorescence are then measured in a plate reader. By comparing eYFP fluorescence in the samples expressing the proteins of interest with eYFP fluorescence in the mCh-P2A-mCh reference construct, novel candidate factors influencing UPR signaling can be identified.

### Reporter optimization

To establish the assay presented in Fig. 1, several conditions were tested and optimized to guarantee the reliability of the assay. First, a suitable UPR responsive promoter had to be identified which shows sufficient strength and high reproducibility in its response to UPR stress. We cloned the promoter regions from four genes that had been reported to be upregulated in ER stress conditions: S-phase kinase-associated protein 1 (SKP1; LOC107761682), bZIP60 (LOC109230966), BIP1 (AT5G28540), and BIP3 (AT1G09080; Iwata & Koizumi, 2005; Ye et al., 2013). These promoters were cloned into plant destination vectors regulating the expression of eYFP, electroporated into *A. tumefaciens*, and infiltrated into *N. benthamiana* leaves. Two days later, we infiltrated the same leaves with 5 μg/mL Tm to induce UPR and measured eYFP levels approximately 24 hours after the second infiltration (Fig. 2A). The regulatory region of SKP1 was the only one that did not lead to a significant increase in eYFP fluorescence after UPR induction. From the remaining promoters, bZIP60 showed the highest fold change of eYFP expression under ER stress conditions (6.03 ± 2.41), followed by BIP1 (5.57 ± 2.19), and BIP3 (4.27 ± 3.51). Considering the high variability observed for pBIP3 and the low fluorescence levels in samples with the bZIP60 promoter, we concluded that pBIP1::eYFP was the most suitable construct for this method. Therefore, all remaining optimization steps were performed using pBIP1::eYFP as the reporter construct.

**Fig 2.**
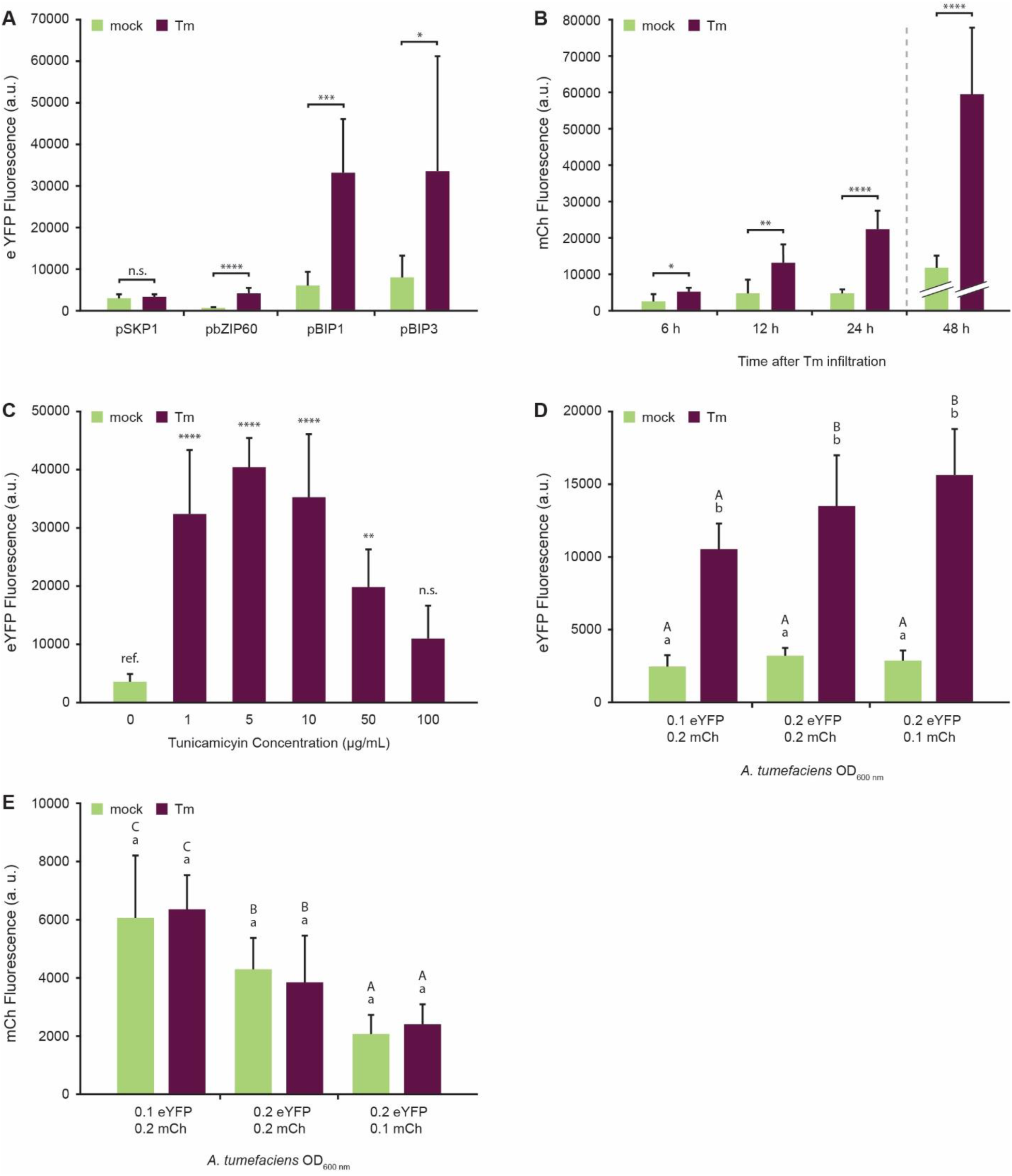
Reporter choice and optimization of unfolded protein response (UPR) induction **(A)** Four reporter constructs were tested for enhanced Yellow Fluorescent Protein (eYFP) upregulation after UPR induction by 5 μg/mL tunicamycin (Tm) infiltration. Subsequent tests were carried out using the pBIP1::eYFP construct. **(B)** Sampling at 6, 12, 24, and 48 hours after Tm infiltration was tested. In the 48 h samples, the gain value of the fluorescence detector was lowered from 100 to 90. The 24 h timepoint was used for further tests. **(C)** Different Tm concentrations were tested for UPR induction. All future tests were done with 5 μg/mL Tm. **(D)** Three optical density at 600 nm (OD_600 nm_) ratios of *A. tumefaciens* strains were tested for optimal eYFP induction after Tm infiltration, and **(E)** candidate protein expression levels, using mCherry (mCh) as a reference. In both **(D)** and **(E)**, the *A. tumefaciens* strain carrying the pBIP1::eYFP reporter construct was co-infiltrated with an *A. tumefaciens* strain carrying a p35S::mCh-P2A-mCh control construct. Error bars represent standard deviation. Error bars represent standard deviation, asterisks represent statistically significant differences (one-way ANOVA or t-test) between samples: * P≤ 0.05, ** P≤ 0.01, *** P≤ 0.001, **** P≤ 0.0001, and n.s. – not significant. Lower case letters represent differences between treatments among samples infiltrated with the same *A. tumefaciens* suspension, while capital letters represent differences within the same treatment among samples infiltrated with different *A. tumefaciens* suspensions in a two-way ANOVA test (P≤ 0.05). a.u. – arbitrary units.

The second factor we optimized was the measurement time after UPR induction. We tested samples at 6, 12, 24, and 48 hours after 5 μg/mL Tm infiltration and compared them to the mock treated samples (Fig. 2B). The timeseries shows a gradual increase in eYFP fluorescence after UPR induction, with the 48 hour timepoint showing overwhelming eYFP levels. In fact, the gain of the detector had to be reduced from 100 to 90 in order to avoid overflow of the signal in these samples, making the arbitrary fluorescence units not directly comparable to the earlier timepoints. However, by comparing the fluorescence fold change between mock and Tm treated plants, we established that there was no further relative induction of promoter activity between the 24 (5.11 ± 1.17) and 48 (5.14 ± 1.57) hour time point. Due to the lower variability in samples measured 24 hours after UPR induction, we decided to use this timepoint in all subsequent experiments.

After determining that the regulatory region of BIP1 displayed a good signal to noise ratio after 24 h of ER stress, we determined the optimal Tm concentration to induce promoter activity. By infiltrating different Tm concentrations in plants transiently expressing eYFP under regulation of the BIP1 promoter, we observed the highest eYFP fluorescence and lowest variation with 5 μg/mL Tm (Fig. 2C). Therefore, this concentration was used for all remaining experiments.

Next, we tested the influence of the ratio between the p35S::mCh-P2A-mCh expression construct and the pBIP1::eYFP reporter vector. Fig. 2D shows the influence of different optical densities at 600 nm (OD_600 nm_) culture ratios in eYFP expression upon ER stress induction. A 1:2 ratio of for pBIP1::eYFP to p35S::mCh-P2A-mCh (OD_600 nm_ = 0.1 and 0.2, respectively) showed the lowest eYFP expression induction. When compared to the other samples however, it showed a similar fluorescence fold change and lower variation (4.40 ± 0.79). An equal ratio of both plasmids (OD_600 nm_ = 0.2) resulted in a 4.28 ± 1.52 fold change, while a 2:1 ratio of pBIP1::eYFP to p35S::mCh-P2A-mCh (OD_600 nm_ = 0.2 and 0.1, respectively) led to a 5.47 ± 1.27 fluorescence increase. Importantly, samples in which the reporter plasmid had a lower OD_600 nm_ relative to the expression plasmid had significantly higher mCh fluorescence (Fig. 2E). Thus, a 1:2 ratio of pBIP1::eYFP to p35S::mCh-P2A-mCh (OD_600 nm_ = 0.1 and 0.2, respectively) leads to similar eYFP induction, while allowing for higher expression of candidate genes. It is also important to note that eYFP induction upon UPR was lower in this assay when compared to the previous experiments. This is likely due to competition in the transient production of two proteins as opposed to one. Nonetheless, in these conditions, eYFP is more than four times more abundant in ER stressed plant leaves.

### Confirmation of UPR induction and proof of principle

To confirm that the assay conditions tested in Fig. 2 and reporter fluorescence correlated with UPR onset, we measured the expression of marker genes by qRT-PCR (Fig. 3A). To that end, the control p35S::mCh-P2A-mCh construct was co-expressed with the pBIP1::eYFP reporter plasmid and the expression of bZIP60, CNX1, SKP1, and PR1 (Chen & Brandizzi, 2013; Hamorsky et al., 2015; Shen et al., 2017; Ye et al., 2011) were measured in both mock and Tm infiltrated leaves. Three of the four marker genes showed a statistically significant upregulation after Tm-induced UPR. In the case of PR1, there seems to be higher expression in UPR conditions but the variability in the dataset and low sample numbers likely led to the observed lack of statistical significance. Nonetheless, this more traditional qRT-PCR based UPR measurement confirmed that the conditions we optimized for our fluorescence-based method leads to ER stress.

**Fig. 3.**
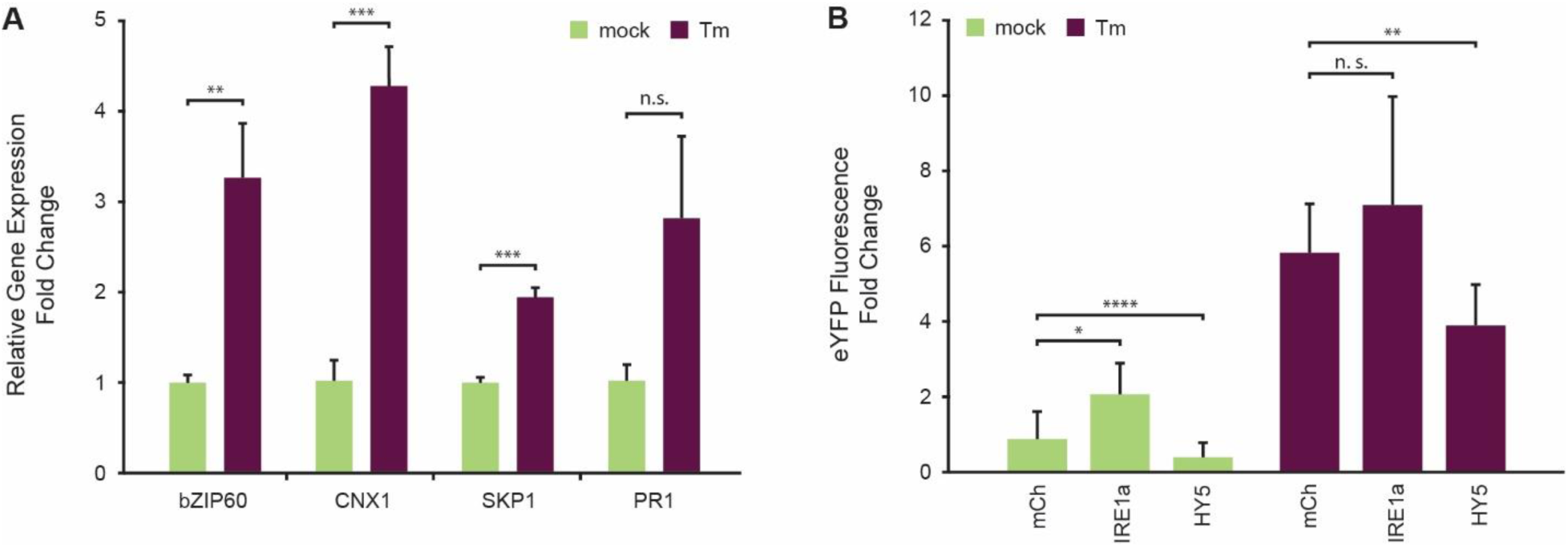
Proof of principle. **(A)** Conditions determined to be optimal for enhanced Yellow Fluorescent Protein (eYFP) upregulation upon tunicamycin (Tm) infiltration were confirmed by measuring unfolded protein response (UPR) marker genes by qRT-PCR. **(B)** Reporter construct expression after co-infiltration with either mCherry (mCh), a UPR signaling component, IRE1a, or the UPR signaling inhibitor, HY5. In both cases, eYFP fluorescence was measured with and without Tm treatment. Error bars represent standard deviation, asterisks represent statistically significant differences (one-way ANOVA or t-test) between samples: * P≤ 0.05, ** P≤ 0.01, and *** P≤ 0.001, when statistical analysis was performed. n.s. – not significant.

Finally, we tested whether our conditions can detect UPR interference using proteins known to be involved in UPR signaling. We co-infiltrated the pBIP1::eYFP reporter construct with either: p35S::mCh-P2A-mCh, as a reference for unaltered UPR signaling; p35S::IRE1a (AT2G17520), which leads to the upregulation of UPR-related genes; or p35S::HY5 (AT5G11260), which is involved in the downregulation of ER stress genes (Fig. 3B; Iwata & Koizumi, 2005; Koizumi et al., 2001; Nawkar et al., 2017). In mock treated samples, we saw a significant induction of eYFP expression caused by the overexpression of IRE1a, showing that this method is capable of identifying proteins that induce UPR signaling. Co-infiltration of Elongated Hypocotyl 5 (HY5) led to a reduction in eYFP upregulation in both mock and Tm treated samples. Taken together, these data show that our method provides a good resolution for identifying proteins that interfere with UPR in plants.

### Library screen and new UPR-interfering protein identification

After optimizing the method with proteins known to have a role in UPR, we aimed to identify novel proteins involved in ER stress signaling. Recent studies showed that some pathogenic effectors can interfere with plant UPR (Fan et al., 2018; Jing et al., 2016). We used a subset of 35 proteins from a library of putative effectors from the biotrophic fungal pathogen *U. maydis* to test whether our method could link any of them to UPR signaling (Fig. 4). In both mock and Tm treated samples, we observed relatively high eYFP fluorescence variation between samples. We therefore decided to apply a strict significance threshold of p ≤ 0.01 in our ANOVA tests. In DMSO (mock) infiltrated plants, only the expression of UMAG_02826_23-399_ – a putative effector expressed without its signal peptide – led to highly significantly increased eYFP fluorescence in *N. benthamiana* cells (Fig. 4A). On the other hand, under ER stress conditions, six putative effectors downregulated eYFP expression, four of which were highly significantly different from the mCh control (p ≤ 0.001; Fig. 4B).

**Fig 4.**
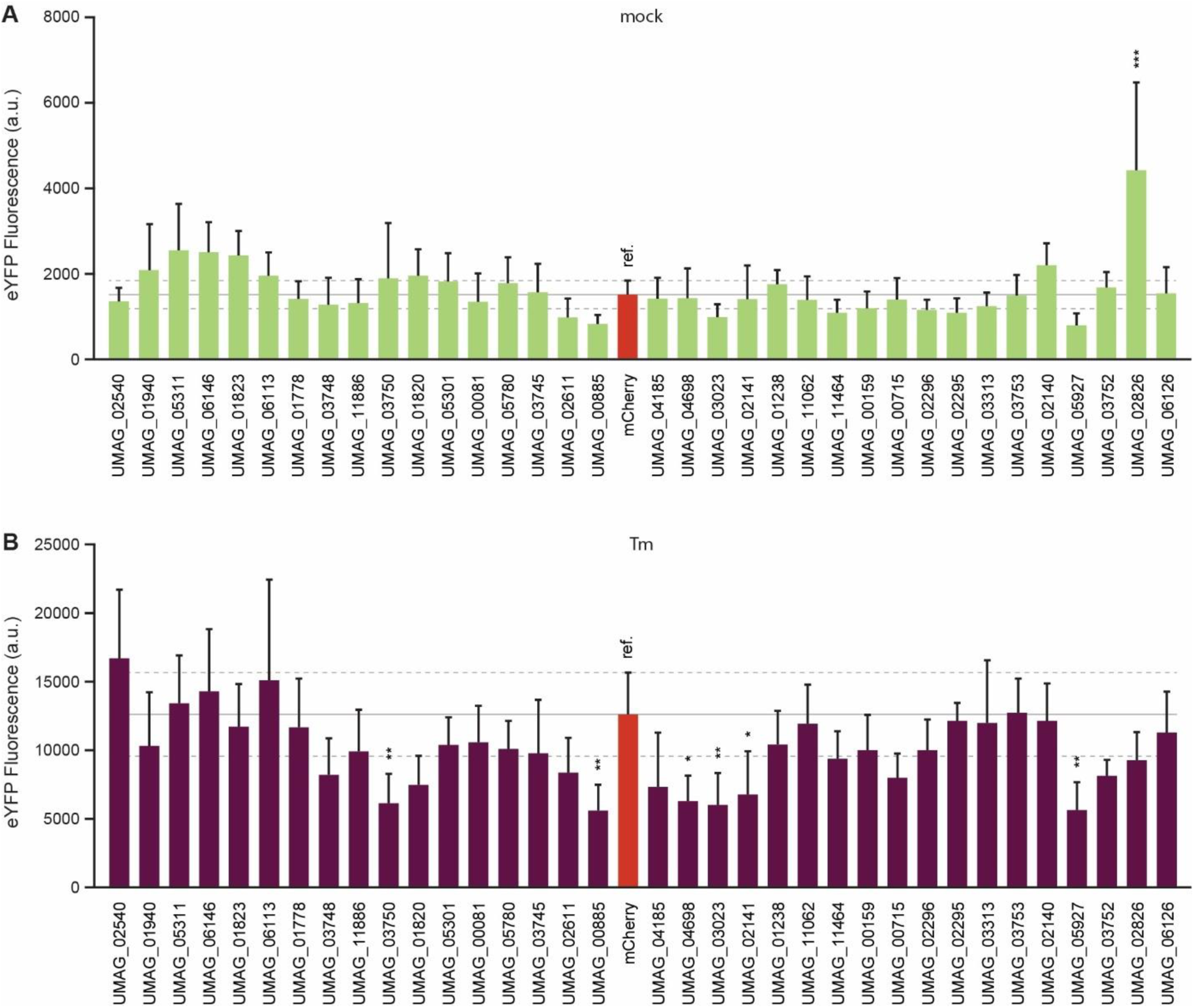
Pilot screen to identify proteins that influence in unfolded protein response (UPR) signaling using a subset of an effector library from the biotrophic plant pathogen *U. maydis*. enhanced Yellow Fluorescent Protein (eYFP) fluorescence in **(A)** DMSO (mock) treated samples to identify proteins that induce UPR signaling, and in **(B)** samples where ER stress was induced (Tm) to identify proteins that inhibit UPR signaling. In both cases, samples of plants expressing mCherry (mCh) were used as a reference. Grey lines represent the average fluorescence (full line) and standard deviation (dashed lines) in mCh samples. Error bars represent standard deviation, asterisks represent statistically significant differences (one-way ANOVA or t-test) between samples: * P≤ 0.01, ** P≤ 0.001. a.u. – arbitrary units.

To confirm these results, we repeated the fluorescence-based assay on the four putative effectors that showed highly significant downregulation of eYFP expression and UMAG_02826_23-399_, which had the opposite effect. In the DMSO treatment, the fold change of eYFP fluorescence relative to the mCh control was relatively consistent in four of the five effectors retested. However, UMAG_02826_23-399_ which significantly upregulated eYFP expression in the first experiment, showed only a slight tendency towards upregulation that was not significant in the second experiment (Fig. 5A). Similarly, variation between the two repetitions in Tm-treated samples was also observed (Fig. 5B). In trying to understand the source of this variation, we considered whether it could be due to changes in protein expression between the two replicates. Because the plasmids encoding the candidate genes also express mCh in equimolar amounts, we used this protein’s fluorescence as an estimate for protein levels of the different constructs (Fig. 5C). We found that there was indeed variation in protein levels between the two replicates in some samples and this is a factor that should be considered when using this method. Nonetheless, the putative effector UMAG_05927_24-370_ consistently downregulated pBIP1 activity to approximately half of what was measured in the mCh control sample (Fig. 5A and B). In Tm infiltrated leaves, qRT-PCR analysis of the same maker genes measured in Fig. 3A showed that expression of UMAG_05927_24-370_ led to a significant decrease in CNX1, SKP1, and PR1 expression, but not bZIP60 (Fig. 5D). This indicates that UMAG_05927_24-370_ can interfere with UPR, either downstream of bZIP60 or in a signaling pathway-specific manner.

**Fig 5.**
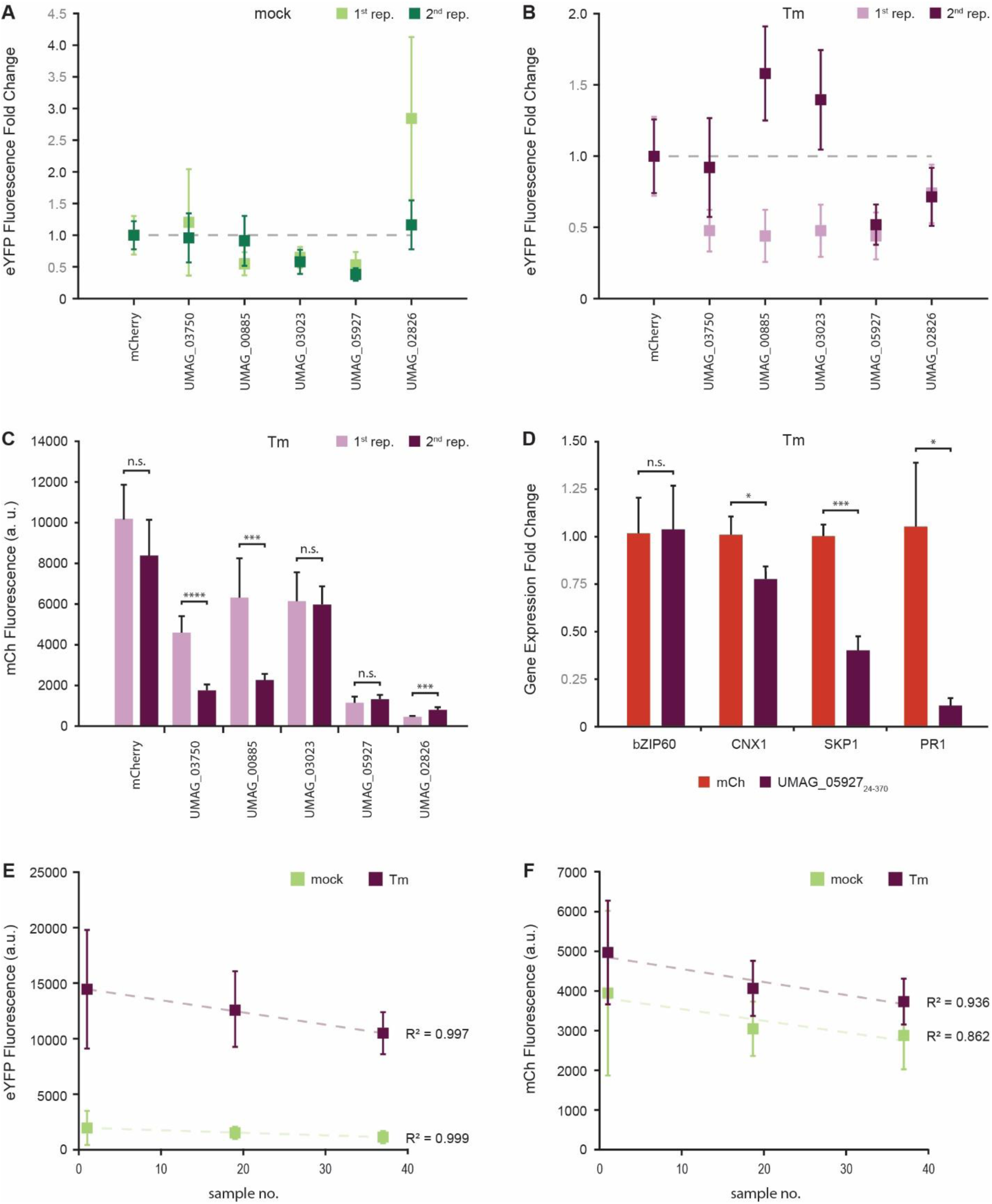
Reproducibility, sources of variability, and confirmation of a protein that interferes with unfolded protein response (UPR) signaling. Variability in enhanced Yellow Fluorescent Protein (eYFP) fluorescence from 2 independent replicates in both **(A)** DMSO (mock) and **(B)** tunicamycin (Tm) treated leaves. **(C)** Estimated variation between replicates of protein expression based on mCherry (mCh) fluorescence in Tm infiltrated samples. **(D)** Relative gene expression of UPR marker genes in samples expressing UMAG_05927_24-370_. **(E)** eYFP and **(F)** mCh fluorescence decrease as a function of sample number. Error bars represent standard deviation, asterisks represent statistically significant differences (one-way ANOVA or t-test) between samples: * * P≤ 0.05, ** P≤ 0.01, and *** P≤ 0.001, when statistical analysis was performed. n.s. – not significant. a.u. – arbitrary units.

There was one more observation we noted that might influence some of the variability of the data. When testing the effect of the 35 putative effectors, we infiltrated the p35S::mCh-P2A-mCh reference construct before, in the middle, and after the infiltration of constructs for effector expression, and measured their fluorescence (Fig. 5E and F). Throughout infiltration, the average signal for both eYFP and mCh fluorescence tend to decrease both in intensity and variability. The only statistically significant decrease was observed in mCh between the first and last samples in Tm infiltrated leaves. Nonetheless, the linear regressions have a high r^2^ fit to the average intensities in all samples. For simplicity, and considering the small scale of our pilot screen, the statistical analysis in Fig. 4 used only the mCh samples from the middle of the assay as a reference. However, if the number of proteins or plants to be tested is increased, a correction factor can be calculated based on the equation from the linear regressions.

## Discussion

UPR is a cellular mechanism that restores homeostasis in stressed cells with highly active transcriptional machineries resulting from abiotic, biotic, or physiological stresses. Due to its importance and ubiquitous nature, the core components that regulate this mechanism are well conserved among eukaryotic organisms and have been characterized in detail (Chakraborty et al., 2016; Iwata & Koizumi, 2012; Strasser, 2018). However, recent studies have been focusing on proteins involved in UPR in specific conditions (Gao et al., 2008; Liu et al., 2007; Meng et al., 2017; Moreno et al., 2012; Nawkar et al., 2017; Pinter et al., 2019; Valente et al., 2009; Xu et al., 2019). This is especially relevant in plants, which rely on signals from their environment to finetune their responses and adapt to diverse changes in their growing conditions. We believe the development of a simple, high-throughput method to identify new factors involved in UPR in plants can lead to important discoveries in this field.

The most commonly used method to link proteins of interest with UPR is qRT-PCR for ER stress marker genes (Chen & Brandizzi, 2013). It requires RNA extraction, cDNA synthesis, and PCR optimization, all prior to experimental testing. This is relatively time-consuming, laborious, expensive, and therefore not suitable to screen libraries of proteins or other molecules. McCormack et al. (2015) described a high-throughput method to screen for the sensitivity of *A. thaliana* to ER stress by growing seedlings in a Tm solution. While this method is simple, efficient, and involves little manipulation of the plant material, its use in identifying new proteins involved in UPR is limited to available seed collections. There are currently no methods available for screening libraries of proteins to identify those that influence UPR in plants.

When studying specific proteins, several studies developed and described small scale methods for specific uses (Hayashi et al., 2013; Liu & Howell, 2010; Meng et al., 2017; Nawkar et al., 2017). While investigating the competition of HY5 with bZip28 for the binding of ER response elements (ERSE), Nawkar et al. (2017) used a construct that upregulated luciferase expression upon ER stress. This enabled them to test the influence of co-expression of two additional proteins on UPR signaling. A similar approach had been described by Iwata & Koizumi (2005) when investigating the regulation of UPR by bZip60 in *A. thaliana*. We have modified and optimized this method to increase its throughput and allow for the simultaneous testing of a large number of proteins for effects on UPR signaling (Fig. 1).

In contrast to other commonly used reporters, fluorescent proteins can be measured directly in leaf discs, leading to minimal sample handling. This results in the reduction of errors that can be introduced in other reporter systems that require further sample preparation steps, such as pipetting inconsistencies, sample mix ups, etc. In addition, fluorescence measurement in leaf discs is fast, reliable, and relatively inexpensive, which dramatically increases the throughput of the method. Furthermore, by using a reference construct, the fold change in eYFP expression can be compared between multiple sampling days and mCh expression can be used as a proxy for transformation efficiency and protein levels. This is achieved by the use of the P2A sequence, which allows for the translation of two separate proteins from the same mRNA molecule in equimolar amounts (Kim et al., 2011). However, the stability of the proteins of interest vary and mCh fluorescence should be used as more of an indicative rather than absolute measure. Nonetheless, antibodies for the P2A peptide are commercially available and a more precise quantification of the proteins can be performed if necessary.

The use of transient protein expression in *N. benthamiana* plants allows for the screening of many candidate genes in a relatively short timeframe, with a restricted growth chamber footprint, and circumvents the restriction of only testing available seed collections. Effectively, this overcomes the gene pool limitations from previous methods, allowing for proteins from virtually any biological source to be screened. However, it has the limitation of restricting the proteins that can potentially be identified to those with conserved targets in *N. benthamiana* UPR signaling. Additionally, inconsistencies in protein expression between samples, as seen in Fig. 5C, E, and F, and known phenotypic changes that occur between transient and stable protein expression have to be taken into account when analyzing data obtained by this method (Bashandy et al., 2015). Because of this, we recommend that an initial screen be used to short list proteins for a second round of testing. Proteins that show a consistent effect on eYFP expression across the two replicates can then be validated by qRT-PCR and further characterized.

Many genes have been reported to be differentially expressed during UPR (Iwata, et al., 2010a; Srivastava et al., 2018). Typically, conserved genes involved in UPR signaling have a basal expression level in most tissues and show a rapid upregulation upon ER stress. From the genes with that expression profile, we tested the regulatory region of 4 of them: SKP1, bZIP60, BIP1, and BIP3 (Fig 2A). In the case of SKP1, Fig. 3A shows that this gene is only moderately upregulated after Tm infiltration. It was therefore not surprising that we could not detect its upregulation in the fluorescence-based assay. This highlights a disadvantage of this method, namely that it is limited to the discovery of proteins with a strong influence on UPR signaling. BiP proteins are essential for UPR and their expression is tightly regulated during this process. bZip60, on the other hand, has a role in early ER stress signaling events and its mRNA is transcribed in non-stress conditions so that it can be unconventionally spliced during UPR (Iwata & Koizumi, 2005; Nagashima et al., 2011). However, the bZIP60 construct tested in Fig 2A showed relatively low levels of eYFP fluorescence in both mock and Tm treated leaves. While the fluorescence fold change was comparable to the promoters of BIP proteins, we considered that the overall low expression could lead to a higher false discovery rate in the identification of proteins with a role in UPR. Regarding the remaining tested promoters, BIP1 has been described to be expressed in low amounts in non-stress conditions and to be upregulated after Tm treatment. On the other hand, BIP3 was found to only be expressed in ER-stress conditions (Iwata et al., 2010b; Maruyama et al., 2014; Nagashima et al., 2014). Surprisingly, fluorescence levels from the BIP1 and BIP3 promoter constructs in non-stress conditions were similar. This could possibly be due to our use of *A. tumefaciens*, which might lead to a small upregulation of UPR genes or to transcription of genes by the bacterium itself. The latter limitation can be overcome by introducing plant specific introns into the coding sequence of the genes, thus preventing their expression by the bacteria (Vancanneyt et al., 1990). Nonetheless, the promoter region of BIP1 showed a more than 4-fold increase in fluorescence after Tm treatment which was sufficient for further testing and proved to be adequate for the purposes of this method.

Another relevant aspect to consider is the induction of UPR itself. In initial experiments, we tested several factors, such as heat stress, ectopic salicylic acid (SA) application, dithiothreitol (DTT) infiltration, and Tm infiltration (data not shown). From these, DTT and Tm infiltrations were the most effective in inducing UPR, with DTT samples showing higher variability in fluorescence intensity. This was most likely due to changes in the cellular redox state which are known to alter the fluorescence of these reporters (Avezov et al., 2013). Additionally, the changes in the redox balance caused by the infiltration of DTT would lead to cellular responses that were not specific to UPR. Therefore, induction of ER stress by Tm infiltration seems to be the most suitable to induce UPR signaling under the conditions tested. However, it is important to note the highly toxic nature of this chemical (Heifetz et al., 1979; Keller et al., 1979; Takatsuki & Tamura, 1971) and appropriate safety precautions should be followed to avoid any direct physical contact with the Tm solution, especially when infiltrating *N. benthamiana* leaves.

The co-expression of the known UPR inducer IRE1a or inhibitor HY5 with our reporter construct showed the expected correlation with eYFP expression following induction of UPR. Together with the measurement of UPR marker genes by qPCR, Fig. 3 shows that the optimal conditions determined in Fig. 2 effectively lead to UPR and that the method is suitable for discovering new proteins that influence this mechanism.

Our small screen with a set of *U. maydis* effectors (Fig. 4) led to the identification of a protein, UMAG_05927_24-370_, which seems to interfere with this process. This effector consistently led to the down regulation of eYFP expression from the reporter construct (Fig. 5A and B) and 3 out of the 4 measured UPR marker genes (Fig. 5D). It is worth noting that the expression of UMAG_05927_24-370_ did not influence bZIP60 transcription, which is commonly upregulated upon ER stress. It did however strongly downregulate pathogenesis related 1 (PR1) expression, which is widely reported to be upregulated upon SA signaling (Seyfferth & Tsuda, 2014). It is tempting to speculate that the influence of UMAG_05927_24-370_ on UPR may be dependent on SA signaling, rather than a more generic UPR inhibition. However. further functional characterization of this protein is needed to better understand its role in UPR interference and pathogenesis. Nonetheless, our method led to the identification of this protein’s involvement in UPR and provided useful hints on how it might function.

## Conclusions

We developed a simple, reliable, and high-throughput method to identify proteins that interfere with plant UPR. Constructs encoding proteins of interest are co-transformed in *N. benthamiana* plants with a fluorescent UPR reporter. Fluorescence is then measured in leaf discs and by comparing control plants with those expressing the protein of interest, in mock or Tm treated samples, that protein’s influence on UPR signaling can be assessed.

Our method enables the testing of gene, and potentially small molecule, libraries using relatively limited resources and time. By using fluorescence as the output of the assay, which can be measured from leaf discs in 96 well plates, many factors can be easily tested in parallel. In fact, our pilot experiment tested 35 proteins and identified one which influences UPR signaling. We anticipate that this reporter system will lead to the discovery of new players in plant UPR signaling, particularly those involved in biotic interactions or that play a role in specific environmental conditions. This will lead to a better understanding of this ubiquitous and very complex cellular homeostasis mechanism and its role in plant biology.

## Methods

### Plant growth conditions

*Nicotiana benthamiana* plants were grown on a 4:1 soil:perlite mixture, at 21°C, 60% humidity and with an 8/16 h dark/light photoperiod in a controlled environment growth chamber. Throughout the growth period, the plants were watered twice per week. *Arabidopsis thaliana* plants for genomic DNA isolation were grown under the same conditions.

### Genomic DNA isolation

Plant genomic DNA for promoter and gene cloning was isolated from leaves of 5 week old plants that were snap-frozen in liquid nitrogen and ground using a Mixer Mill MM 400 (Retsch GmbH, Germany) for 1 min 30 sec at 30 Hz. To the resulting powder, 500 μL of extraction buffer (5.5 M Guanidine Thiocyanate, 20 nM Tris-HCl, pH 6.6) was added and the sample was vigorously vortexed before centrifugation at 20,000 x *g* for 5 min. The supernatant was loaded into an EconoSpin® All-In-One Silica Membrane Mini Spin Column (Epoch Life Science, INC., USA) and centrifuged at 20,000 x g for 1 min. The membranes were washed twice with cleaning buffer (80% ethanol, 10 mM Tris-HCl, pH 7.5) and centrifuged at 20,000 g for 1 min. The DNA was eluted with 50 μL of purified water and stored at -20°C until further use.

### Vector construction

DNA manipulation and plasmid assembly were performed according to standard molecular cloning procedures (Ausubel et al.,1987; Sambrook & Russell, 2006), using the GreenGate vector set and cloning conditions (Lampropoulos et al., 2013). All DNA manipulations were performed using the *Escherichia coli* MACH1 strain (Thermo Fisher Scientific, USA). Cloned genes and promoter sequences were blunt-end ligated into the pJet vector (Thermo Fisher Scientific, Waltham, MA, USA) before further golden gate cloning procedures. The plasmids used from the GreenGate vector set have the following Addgene IDs:48815, 48820, 48828, 48834, 48841, 48848, and 48868. Primers used in this study are listed in Table 1. Whenever necessary, BsaI restriction sites native to the coding sequences of the promoters or putative effectors were mutated. Silent mutations were introduced by site directed mutagenesis (Liu & Naismith, 2008) to preserve the native amino acid sequence and maintain the efficiency of the Golden Gate cloning method (Engler et al., 2008). In the case of the fluorophores, eYFP was re-cloned from a different vector system using primers with adaptors to enable its compatibility with our cloning strategy (Table 1). Nested PCR from the Addgene vector 48828 was performed to create the P2A-lifeact-mCh CD module compatible with the GreenGate vector set (forward primer 1 – atatggtctcatcagctGGTTCTGGAGCTACTAACTTCTCTCTCTTGAAGCAAGCAGGAGATGTGGAAGAAAACCCTGGTCCAATG, forward primer 2 – AAGAAAACCCTGGTCCAATGGGTGTCGCAGATTTGATCAAGAAATTCGAAAGCATCTCAAAGGAAGAAGTGAGCAAGGGCGAGGA, and reverse primer – atatggtctctgcagctaCTTGTACAGCTCGTCCA). The lifeact sequence, which attaches the fluorophore to actin filaments, was originally planned for the effector library used here. Because mCh fluorescence was merely used for estimating protein expression, we refer to this part of the construct as “P2A-mCh” for simplicity.

**Table 1.**
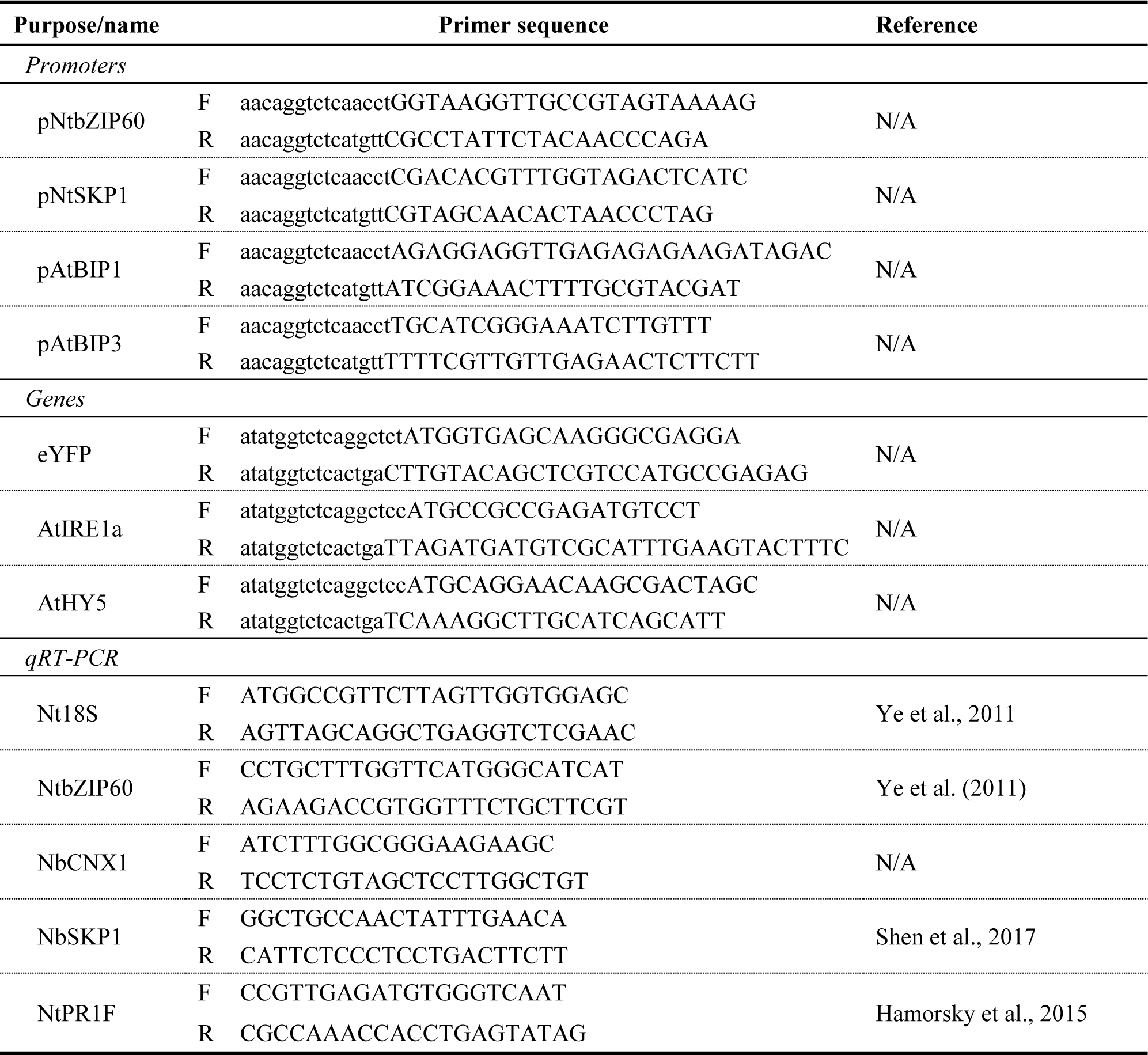
List of primers used for promoter and gene isolation, and relative gene expression measurement by qRT-PCR. Small letters in the primer sequence represent adapters for golden gate cloning, compatible with the GreenGate vector set (Lampropoulos et al., 2013). F and R represent forward and reverse primer sequences, respectively.

The library of putative effectors was cloned based on the effector prediction analysis described in Mueller et al., 2008. Genes, specific primer sequences used to isolate them, and the updated signal peptide prediction scores calculated in SignalP v5.0 (Armenteros et al., 2019) was recently described in Alcântara et al. (2019). All putative effectors were cloned without the predicted signal peptide.

### *Agrobacterium tumefaciens* infiltration and UPR induction

Plasmids were transformed into *A. tumefaciens* strain GV3101 (pSoup) by electroporation (Holsters et al., 1980; Lampropoulos et al., 2013) Transformed cells were selected on Luria broth (LB)-agar media supplemented with antibiotics (50 μg/mL rifampicin, 100 μg/mL spectinomycin, 50 μg/mL gentamycin) and grown at 28°C for 2 days. Colonies were then grown overnight in liquid LB medium supplemented with the same antibiotics, 20 μM acetosyringone, and 10 mM 2-(N-morpholino)ethanesulfonic acid (MES, pH 5.6). When necessary, glycerol stocks of the strains in liquid culture were done by adding glycerol to a final concentration of 40% v/v and freezing at -80°C until further use. Liquid cultures were pelleted at 3000 x g for 10 min and resuspended in 10 mM MES, pH 5.6, 10 mM Magnesium chloride, and 0.15 mM acetosyringone. OD_600 nm_ was measured and the cultures were diluted and mixed with the strain carrying the reporter construct to the final target OD_600 nm_. The suspensions were then left at room temperature for a minimum of 3 hours to allow for the expression of virulence genes. Finally, each bacterial mixture was co-infiltrated in the first two fully developed leaves from two tobacco plants (4 leaves/suspension in total). After 2 days, either DMSO (mock treatment) or tunicamycin (Tm; UPR induction) were infiltrated into the same leaves. Tm stock solutions were dissolved in DMSO to a concentration of 1 mg/mL and frozen at -20°C until further use. Mock treatments were typically infiltration of a 0.5 % DMSO solution, the same as the final 5 μg/mL Tm solution.

### Fluorescence measurements

One day after the second infiltration step, four discs from each infiltrated leaf were collected with a disposable 4 mm biopsy punch (Integra York PA, Inc, USA), and floated on 100 μL of water in 96 well black plates. Leaf disc fluorescence was measured in a Synergy H1 Hybrid Multi-Mode Microplate Reader (BioTek Instruments, Inc, USA). eYFP was excited at 485 nm and measured at 528 nm, while mCh was excited at 570 nm and measured at 610 nm. Autofluorescence was measured in uninfiltrated leaves and the averaged value was subtracted from all fluorescence measurements.

### Quantitative real-time polymerase chain reaction (qRT-PCR)

qRT-PCR was performed as described in Rabe et al. (2016). Briefly, RNA was extracted from infiltrated tobacco leaves in 3 independent replicates, using the RNeasy Plant Mini Kit following the manufacturer’s protocol (QIAGEN Inc., Germantown, MD, USA). DNA was removed with the RapidOut DNA Removal Kit, and reverse transcription was performed using the RevertAid H Minus First Strand cDNA Synthesis Kit (Thermo Fisher Scientific, Waltham, MA, USA). qRT-PCR measurements were performed with the Roche LightCycler® 96 system according to manufacturer’s instructions (Roche Diagnostics, Rotkreuz, Switzerland). Relative expression values were calculated by the 2^-ΔΔCt^ method (Livak & Schmittgen, 2001). All primers used are listed in Table 1.

### Statistical analysis

Statistical significance was tested in GraphPad Prism 8.0.2 (2019). T-tests, one-way or two-way analysis of variance (ANOVA) followed by a multiple comparison Tukey hypothesis testing were used when appropriate. In each sample, two leaves of two plants were infiltrated twice and each infiltration spot (8 in total per sample) was considered a technical replicate.

## List of abbreviations

BiP: luminol binding protein;
bZIP: basic leucin zipper;
DTT: dithiothreitol;
ER: endoplasmic reticulum;
ERSE: ER stress elements;
HY5: Elongated Hypocotyl 5;
IRE1: Inositol-requiring enzyme 1;
LB: Luria broth cell culture medium;
mCh: mCherry;
MES: 2-(N-morpholino)ethanesulfonic acid;
OD_600 nm_: optical density at 600 nm;
p35S: CaMV 35S promoter;
PCD: programmed cell death;
qRT-PCR: quantitative real time polymerase chain reaction;
S1P: site 1 protease;
S2P: site 2 protease;
SA: Salicylic acid;
Skp1: S-phase kinase-associated protein 1;
Tm: tunicamycin;
UPR: unfolded protein response;
UPRE: UPR responsive elements;
eYFP: enhanced yellow fluorescent protein.

## Author contributions

AA designed the study; AA, DS, and FN did the experimental work; AA analyzed the data and wrote the manuscript. AD supervised the project and provided critical feedback. All authors read and approved the final manuscript.

## Acknowledgements

We would like to acknowledge the GMI/IMBA/IMP service facilities, particularly the molecular biology services for Sanger sequencing and support when using the plate reader. We would also like to thank the Plant Sciences Facility at Vienna BioCenter Core Facilities GmbH (VBCF), member of the Vienna BioCenter (VBC) for providing the use of their plant growth facilities. We would also like to acknowledge Dr. J. Matthew Watson for input on the manuscript.

## Availability of data and materials

Vectors and vector maps containing detailed sequence information necessary to use this method are available from Addgene (Massachussetts, USA; Table 2). Detailed information on the remaining plasmids is available upon request.

**Table 2.**
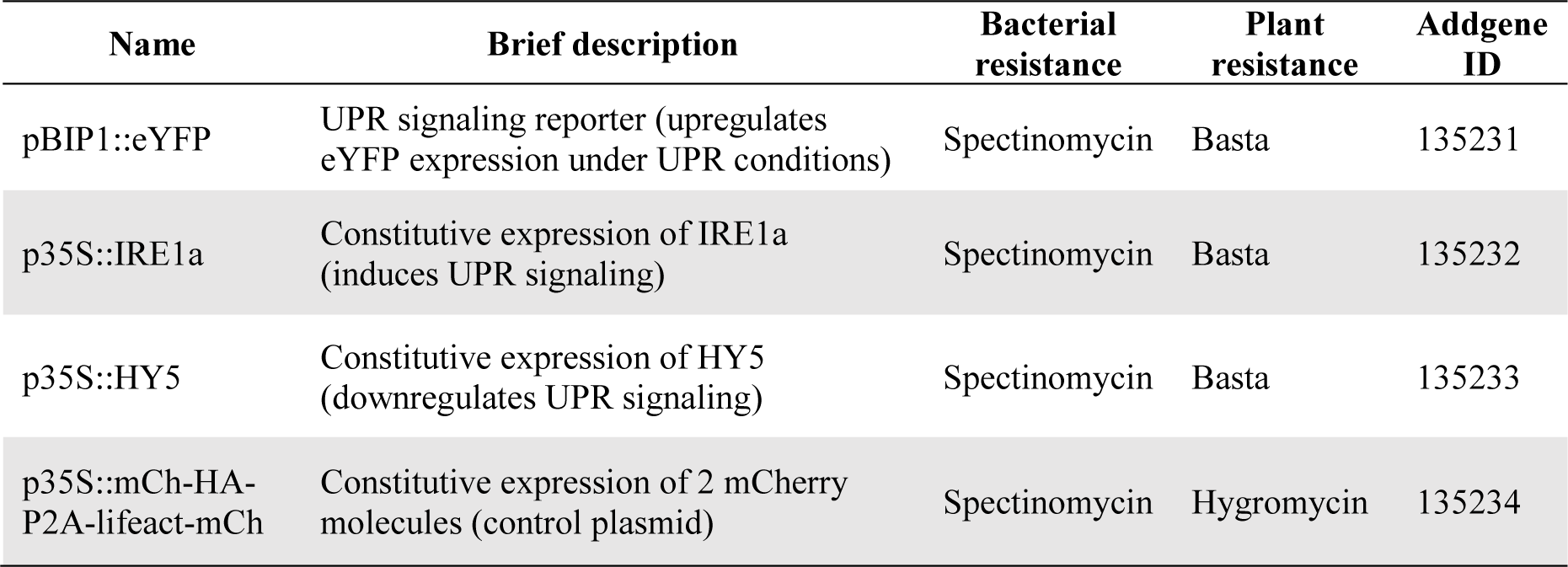
Publicly available vector set to use this method.

